# COMBINe: Automated Detection and Classification of Neurons and Astrocytes in Tissue Cleared Mouse Brains

**DOI:** 10.1101/2022.12.06.519291

**Authors:** Yuheng Cai, Xuying Zhang, Chen Li, H. Troy Ghashghaei, Alon Greenbaum

## Abstract

Tissue clearing renders entire organs transparent to enable combination with light sheet fluorescence microscopy and accelerate whole tissue imaging. Yet, challenges remain in analyzing the large resulting 3D datasets that consist of terabytes of images and information on millions of labeled cells. Previous work has established pipelines for automated analysis of tissue cleared mouse brains. However, they have focused on single color channels and/or detection of nuclear localized signals, in relatively low-resolution images. To address this gap, we present an automated workflow to map labeled neurons and astrocytes in the genetically distinct Mosaic Analysis with Double Markers (MADM) mouse forebrains. We named the workflow COMBINe (Cell detectiOn in Mouse BraIN) as it combines modules from multiple pipelines. With RetinaNet in its core, we quantitatively analyzed the regional and subregional effects of MADM-based deletion of the Epidermal growth factor receptor on neuronal and astrocyte populations in the mouse forebrain.

## Motivation

Tissue clearing has been extensively used to explore cellular functioning of entire organs. By imaging intact mouse brains with cellular resolution, the resulting datasets contain terabytes of images to process and millions of labeled cells that need to be counted and classified based on morphology and color. Therefore, automation of data processing and analysis is in urgent need. We presented a workflow (COMBINe) to automatically locate and classify labeled cells in such 3D datasets and to quantitatively analyze the regional effects after registration to the referenced Allen Brain Atlas. We applied the approach to study the effects of sparse Epidermal growth factor receptor deletion in glial progeny.

## Introduction

Recent advances in tissue clearing methods have enabled three-dimensional (3D) imaging of intact mammalian organs and entire small organisms while preserving comprehensive structural and cellular information (R. Cai et al., 2019; Chung & Deisseroth, 2013; Ertürk et al., 2012; Nudell et al., 2022; Renier et al., 2016; Richardson & Lichtman, 2015; Susaki et al., 2015). Using light sheet fluorescence microscopy (LSFM), high-speed imaging of cleared and immunolabeled tissues can be achieved (Kubota et al., 2017; Moatti et al., 2022; Ueda et al., 2020; Zhao et al., 2020). Yet, challenges remain in analyzing the large resulting 3D datasets with terabytes of images and information on millions of labeled cells. Hence, the automation of data processing and cell detection is paramount for high throughput cellular profiling of cleared tissues.

Previous work has established pipelines for automated analysis of tissue cleared mouse brains such as ClearMap (Renier et al., 2016), CUBIC (Matsumoto et al., 2019) and NuMorph (Krupa et al., 2021). ClearMap presents an immunolabeling-enabled tissue clearing method with superior efficiency in data processing, accomplishing automated cell counting, dataset registration, and statistical analysis. The datasets are analyzed using traditional image processing routines (e.g., morphological operations). CUBIC incorporates improved clearing, imaging, and cell-nucleus-detection protocols, with the ability to quantify cells of entire adult mouse brains. NuMorph is an advanced end-to-end data processing tool for accurate cell-type quantification within the mouse cortex, which reaches high precision in nuclei detection. The previous pipelines achieved high efficacy, yet they focused on single color channels and/or detection of nuclear signals, in relatively low-resolution images. As such, the existing pipelines fail to capture cases where 1) whole cells or cell membranes rather than nuclei are stained, 2) colocalization of color channels is crucial, and 3) the cells can be classified based on their morphology, which is obtainable at high-resolution. These requirements are paramount broadly in neuroscience research, especially those that require in situ analysis of multiple genetic markers as in Confetti mice (Zhou et al., 2019) or Mosaic Analysis with Double Markers (MADM; Beattie et al., 2017; Johnson & Ghashghaei, 2020; Liang et al., 2012, 2013; Zhang et al., 2020). Here we addressed this critical gap by building on the strengths of existing pipelines while developing a new automated workflow that was tested in MADM datasets. In these datasets the fluorescent expression in neurons and glia report on their distinct genotypes.

In the core of our workflow is deep learning, a data driven approach. Recent adaptation of deep learning (LeCun et al., 2015) in biomedical studies has enabled automated and accelerated image processing (Bayramoglu et al., 2016; Hollandi et al., 2020; Sità et al., 2022). In combination with tissue clearing and 3D microscopy, deep learning has also been utilized to study cancer metastasis in mice (Pan et al., 2019), segment mouse brain vasculature (Todorov et al., 2020), and detect crown-like structures in adipose tissues (Geng et al., 2021). Inspired with these studies, we have surveyed the deep learning literature, and employ a deep learning model (RetinaNet) that show superior performance in dense object detection and apply it to a neurodevelopmental problem (Y. Cai et al., 2021; Lin et al., 2018; Waithe et al., 2020).

With RetinaNet in its core, we developed an automated workflow we refer to as COMBINe to map labeled neurons and astrocytes in the forebrains of three genetically distinct MADM mice (Figure 1). We used a conditional mouse allele for the Epidermal growth factor receptor (*Egfr*) in combination with MADM alleles. Using established *Emx1^cre^* line to restrict recombination to the dorsal telencephalon (cerebral cortices and the hippocampal formation), three genotypes were obtained: *Emx:MADM:*+/+, *Emx:MADM:F*/+, *and Emx:MADM:F/F* (Zhang et al., 2022). We have recently reported on variable effects on specification and differentiation of MADM glia in cerebral cortices of these mice, which has led to the developmental model that EGFR-dependent and EGFR-independent perinatal gliogenesis regulate forebrain development cooperatively (Zhang et al., 2022).

**Figure 1.**
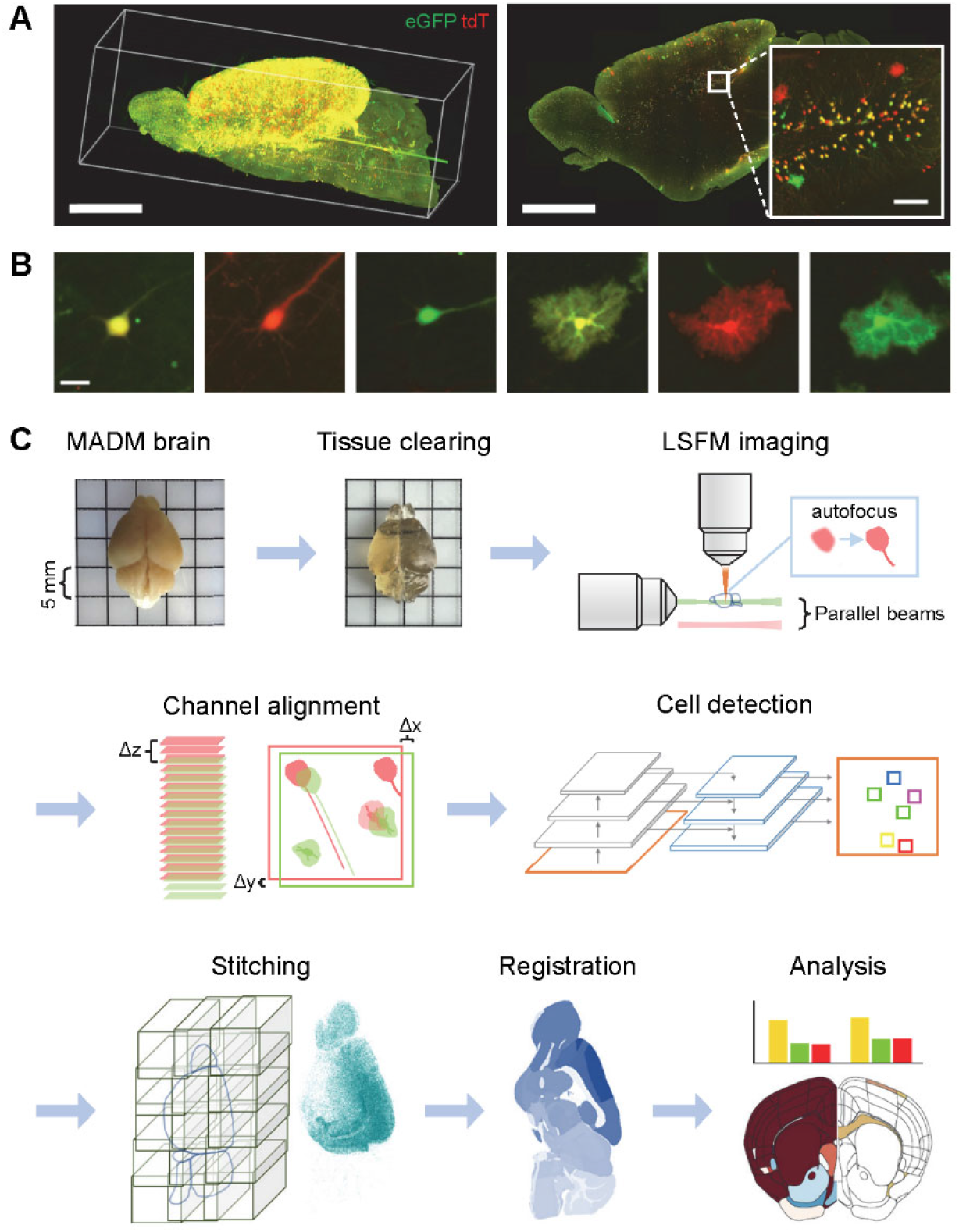
Automated cell detection in the cleared MADM mouse forebrain. **(A)** Left: Maximum intensity projection of fluorescence signals from a hemisphere of an *Emx:MADM:*+/+ forebrain labeled with antibodies to enhance the endogenous eGFP (green) and tdTomato (tdT, red) signals (scale bar, 3 mm). Right: A virtual 30 μm thick slice from the same cleared sample (scale bar, 2.5 mm) with a zoomed-in area (scale bar, 100 μm). eGFP – enhanced green fluorescent protein, tdT – a red fluorescent protein. **(B)** Magnified images of three isolated MADM-labeled neurons and astrocytes expressing eGFP (green), tdT (red), or both (yellow) MADM reporters. Scale bars, 25 μm. **(C)** The proposed COMBINe workflow for MADM cell mapping (see also Figure S1).

Here we used the *Egfr* MADM model to test COMBINe. Using tissue clearing and a custom-built LSFM that changes its properties based on the sample, we generated high-quality MADM datasets with cellular resolutions sufficient to morphologically differentiate astrocytes from neurons. Employing RetinaNet, 0.5 million MADM cells across six classes (yellow, red, green: neurons and astrocytes) were located and counted from each MADM forebrain hemisphere. Taking advantage of available packages, stitched volumes and detected cells were aligned to maps provided by the Allen Brain Atlas (ABA) for regional registration and analysis (Wang et al., 2020). Significant variations in cell populations corresponding to the MADM genotypes were confirmed matching our findings from laborious manual counting and registrations.

## Results

### Automated cell detection in cleared MADM mouse forebrains

MADM allows simultaneous labeling and genetic manipulation in clones of somatic cells in isolated and sparse populations of progenitors (Zong et al., 2005) and has been successfully employed in genotype-phenotype studies on neurogenesis and gliogenesis (Beattie et al., 2017; Hippenmeyer et al., 2010; Johnson & Ghashghaei, 2020; Laukoter et al., 2020; Liang et al., 2013; Zhang et al., 2020). MADM-labeled neurons and glia can possess distinct genotypes, which was tracked by permanent labeling of two mitotically-derived daughter cells with two distinct and nonoverlapping fluorescent markers: enhanced green fluorescent protein (eGFP) and/or tdTomato (tdT; Figure 1A). An entire *Emx:MADM:*+/+ mouse forebrain contained nearly one million labeled cells at postnatal day 30 (P30) which constitutes the early stages of juvenile life in mice. MADM cells in these forebrain samples can be counted and classified into six groups based on morphology and color: red (tdT), green (eGFP) and yellow (both reporters coexpressed) neurons and glia (Figure 1B).

Increased understanding of the complexities in MADM data in the mouse forebrain inspired the need for classification of MADM labeled cells based on morphology. In addition, MADM colors convey information regarding genotype of cells which required colocalization of multiple color channels. Therefore, we designed a workflow that spanned tissue clearing to quantitative analysis (Figure 1C). *Emx:MADM* mouse brains were split into hemispheres, cleared using the iDISCO+ protocol (Renier et al., 2016) and imaged by a custom-built LSFM (Li et al., 2021, 2022; Moatti et al., 2020). The LSFM was equipped with an autofocus algorithm to correct the shift between the light sheet illumination beam and the detection focal plane, and therefore, our images were crisp and focused throughout acquisition. Before imaging, illumination beams of different wavelengths were aligned parallel to ensure accurate data processing (see later). Imaging with subcellular resolution, the acquired two-channel datasets consisted of ~ 400 GB of data per hemisphere. Channel alignment was achieved using NuMorph (Krupa et al., 2021) that locally evaluated the translation between the color channels, followed by a RetinaNet model trained to detect six classes of labeled cells simultaneously (Y. Cai et al., 2021; Lin et al., 2018). The aligned tiles and detected cells were then stitched according to TeraStitcher (Bria & Iannello, 2012) and registered to the ABA by ClearMap (Renier et al., 2016) for regional analysis in the forebrain. By mixing and matching codes from different packages we were able to build on their relative advantages (Figure S1). Taking advantage of the abovementioned tools, we were able to analyze a single *Emx:MADM* hemisphere in four days using a regular desktop computer and employing the proposed pipeline. For completeness, glia labeled using *Emx:MADM* forebrains consisted of oligodendrocytes and astrocytes. However, the morphology and size of mature astrocytes was highly distinguishable, whereas oligodendrocytes were small and difficult to detect, and their numbers were small compared to neurons and astrocytes (Zhang et al., 2022). Thus, we trained our platform to distinguish between neurons and astrocytes.

### COMBINe is highly suitable for automated detection, stitching and registration of MADM cells in the P30 cleared forebrain

RetinaNet was adapted to detect six classes of MADM cells: yellow, red, and green neurons and astrocytes. We selected the RetinaNet model because of its superior performance in dense object detection of real-world images (Zou et al., 2019) and in cell detection of fluorescence microscopy images (Waithe et al., 2020). Cell detection was performed after color registration, as local translation was sufficient to register the color channels under parallel beam condition (Figure 2A; see next section). Cell detection was performed in 2D instead of 3D, due to the difficulty of labeling and training, as well as the variance in resolution along different axes in 3D images. We explored the training regime of the network and applied it to fluorescence images of MADM brain sections in a pervious study (Y. Cai et al., 2021). We again employed the data augmentation of color swap and saturation to balance the color classes and utilized transfer learning to train the network on a pretrained backbone. After training, the model which performed best on validation data was selected as the inference model. The inference model reached an average precision of 0.862 across six classes on unseen test data (Figure 2B). An average precision of 0.897 was achieved for detection of neurons, while an average precision of 0.826 was achieved for detection of astrocytes. The precision-recall curves across six classes are shown in Figure 2C with several examples of the model predications shown in Figure 2D. The color classification was robust even in cases with minor misalignment between the two-color channels (Figure 2E).

**Figure 2.**
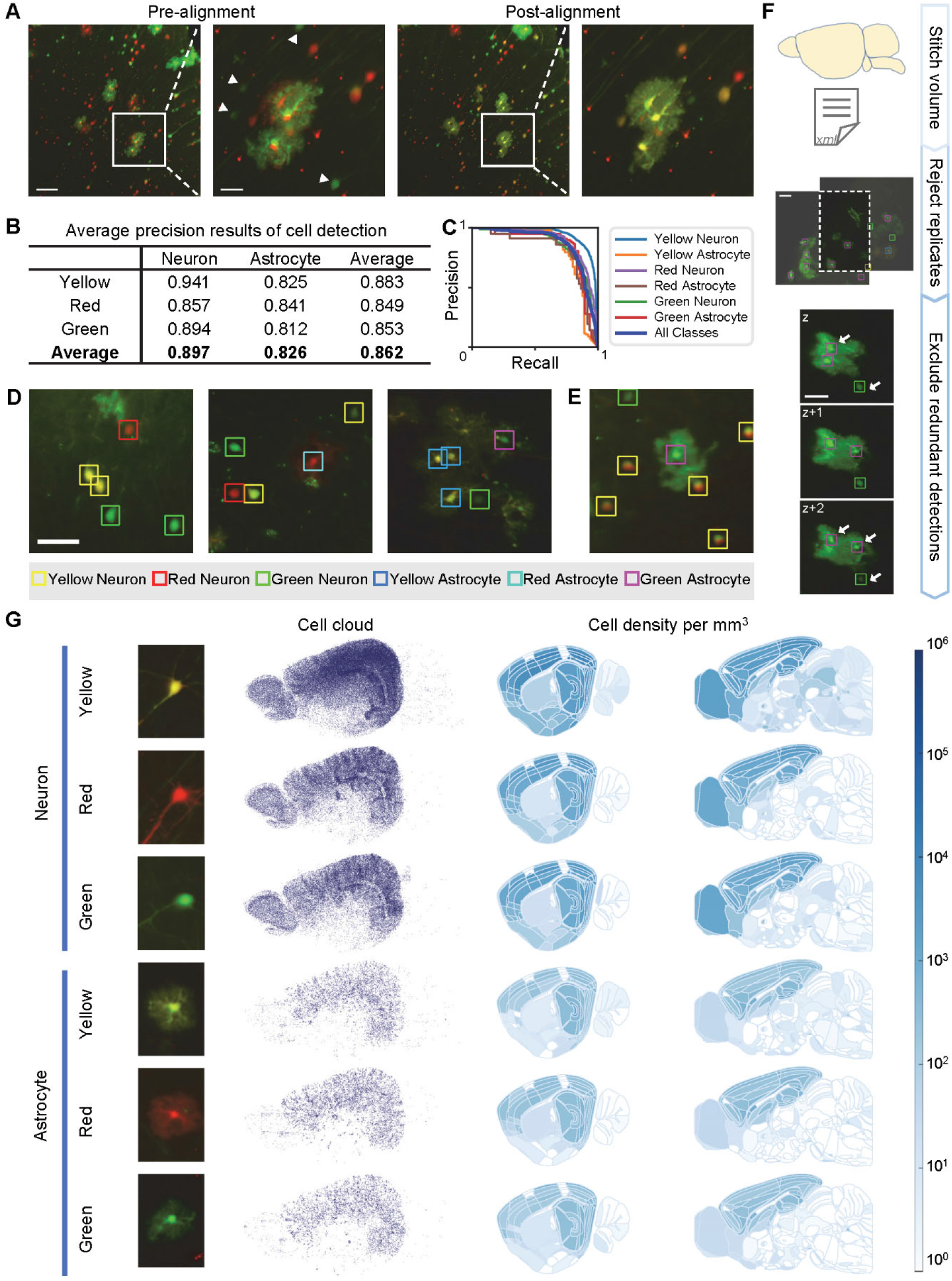
Automated detection, stitching and registration of labeled cells. **(A)** Images pre- and post-channel alignment from parallel illuminations (left panel; scale bar, 100 μm). Cells in the wrong z-plane (arrowheads) disappear following alignment. Boxed areas are zoomed in images (scale bar, 30 μm). **(B)** An overall average precision of 0.86 was achieved across the six cell classes using RetinaNet. **(C)** Precision-recall curves. **(D-E)** Representative images in which the program made predictions with accurate (D) and inaccurate (E) color registrations (scale bar, 50 μm). **(F)** Stitching and merging of detected cells in 3D for each tile were according to TeraStitcher parameters. Replicates in overlapped regions across tiles were rejected, and redundant detections across different z-planes were excluded according to signal intensities. **(G)** Detected cell clouds and corresponding regional densities after registration to the Allen Brain Atlas (ABA), in a single hemisphere of an *Emx:MADM:*+/+ forebrain.

After detecting the cells in raw tiles, the displacement parameters between individual tiles were estimated using TeraStitcher (Bria & Iannello, 2012), but the high-resolution volume was not stitched. A custom script using the displacement parameters was created for excluding duplicate detections in overlapping regions between adjacent tiles and removing redundant detections across different z-planes (Figure 2F; see Methods). Analysis of a cleared MADM hemisphere was accomplished in four days using a standard computer with generated datasets containing approximately 100,000 images per sample (more than 1.6 million image patches for cell detection). More than 400,000 MADM cells were detected in each cleared hemisphere. A low-resolution stitched volume of the hemisphere was later registered to brain regions annotated by ABA using ClearMap. Detected MADM cells and corresponding regional density maps of a representative *Emx:MADM:*+/+ sample are visualized in Figure 2G using a custom script. Nearly all *Emx:MADM:*+/+ cells were in the dorsal forebrain consisting of the olfactory bulb, the cortex, and the hippocampus, which was consistent with previous observation on 2D slices (Liang et al., 2012). As a quality check for our color registration process, the yellow, red, and green cells portion of the total population was approximately 50%, 25% and 25% respectively, which was in line with previous estimates (Hippenmeyer et al., 2010).

In summary COMBINe can detect and classify cells in an entire MADM mouse brain hemisphere. The raw datasets were acquired using our custom adaptive LSFM, and since commercial LSFMs are not adaptive, and do not change their properties on the fly, we have tested the effect of two realistic acquisition scenarios on the performance of the RetinaNet model in the next section: 1) when the cells are slightly out-of-focus, 2) when the illumination beams (one per color) are not perfectly parallel.

### RetinaNet accurately detects out-of-focus cells

LSFM images can seem out-of-focus when the objective focal plane does not perfectly overlap with the illumination beam (Tomer et al., 2015). Therefore, we tested the performance of our network in images containing blurry cells using the same tiles that were acquired in- and out-of-focus. To our surprise, the RetinaNet model performed well in detection and color classification of cells although degradation in image quality was observed in the out-of-focus images. This finding illustrated the robustness of the RetinaNet model, and its ability to generalize well in less-than-optimal conditions (Figure 3A). It is important to note that while training the RetinaNet model, cells from the edges of the field-of-view that exhibited lower contrast in comparison to the center of the field-of-view were included. This phenomenon of fluctuations in image quality as a function of position within the field-of-view was an inherent property of using a gaussian illumination beam. Consequently, the network was trained with variability in image contrast, which may explain the robustness of the network in correctly detecting out-of-focus cells (Figure 3A parallel beam scenario).

**Figure 3.**
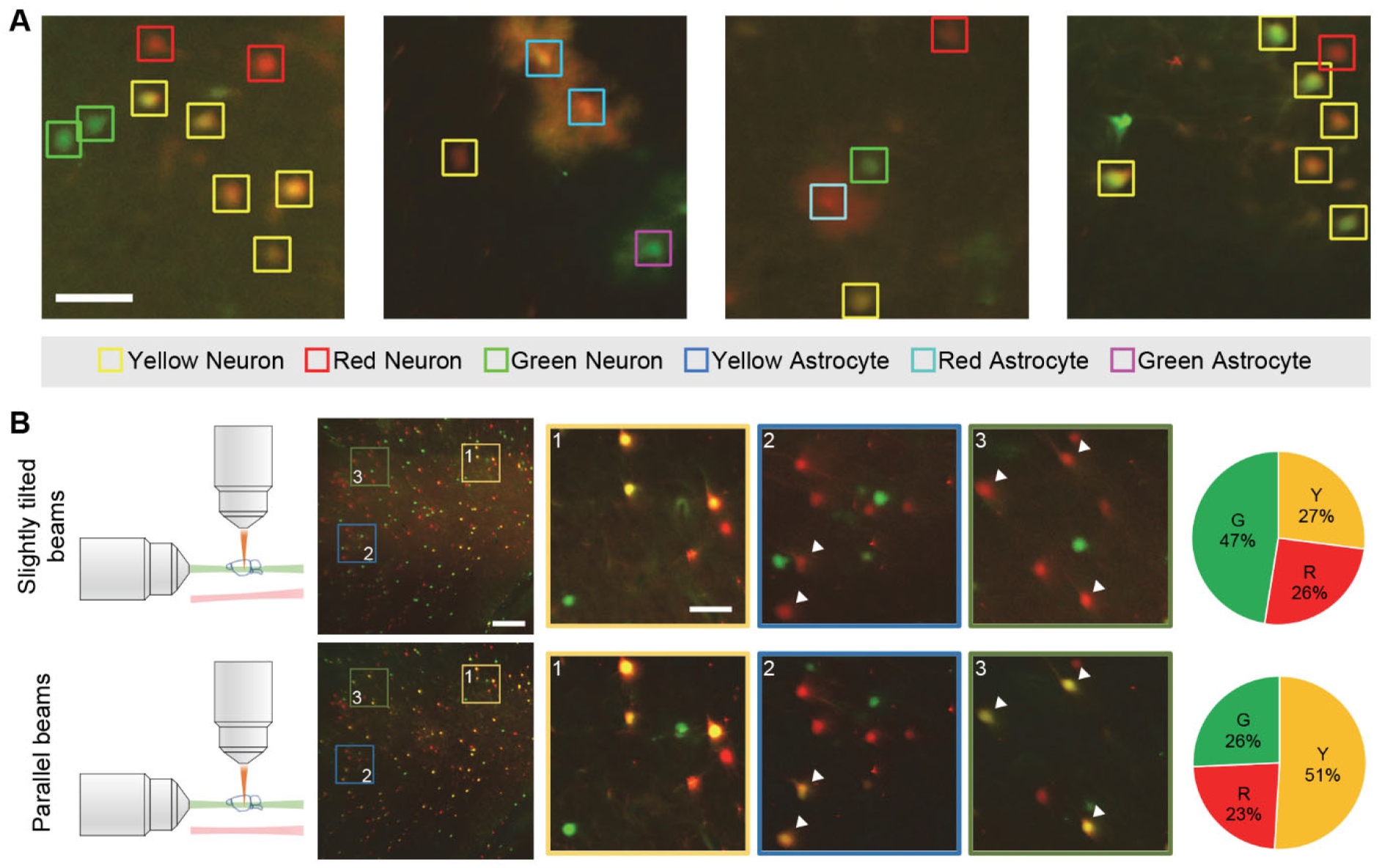
RetinaNet detection is robust for out-of-focus cells. **(A)** Representative predictions in images of out-of-focus cells (scale bar, 50 μm). **(B)** Illustrations of LSFM geometry for tilted (upper) and parallel (bottom) illuminations. Images from the same field-of-view (scale bar, 200 μm) are shown with three zoomed in areas (scale bar, 25 μm). Unparallel beams during imaging result in misalignment between the two channels (e.g., the two-color channels are appropriately overlapped in zoomed region 1 but misaligned in regions 2 and 3 as marked by arrowheads). Erroneous color classification ensues resulting in fewer yellow cells detected (pie charts).

Next, we tested our pipeline in another scenario, when the illumination beams of different color channels propagated through slightly different light paths during LSFM imaging. This can occur when the microscope setup is out of alignment due to angular drift of the scanning galvo regarding to the room temperature, a laser beam shift after long period of imaging (e.g., air bubbles) and more. This situation resulted in only partial overlap of the color channels i.e., misalignments in some regions even after registration of the color channels using NuMorph. Importantly, misalignment of the color channels influenced the subsequent quantitative analyses (see pie charts in Figure 3B). In wild type MADM mice, we expected a color ratio of 50% yellow, 25% red, and 25% green neurons, and we have used this distribution to test the performance of our color registration. This approximate ratio was achieved when the illumination beams were parallel to each other, and the different color channels were registered using local x-y-z translation. However, with slightly tilted illumination beams this ratio was greatly distorted. We observed fewer yellow cells and more green cells.

Together these findings suggest that the pipeline and the RetinaNet model were robust and highly effective for detection and color classification, even in cases containing unfocused images, or slight misalignments between the color channels, as long as the light sheet illumination beams were parallel to each other.

### COMBINe faithfully captures and detects elevation of red (wildtype) astrocytes and depletion of green (Egfr-null) astrocytes in the Emx:MADM:F/+ forebrain

We tested COMBINe to compare distinct MADM datasets in detail (Beattie et al., 2017; Zhang et al., 2020) in which *Emx1 driven* cre recombinase expression (*Emx1^cre^*) is restricted to the dorsal telencephalon (Liang et al., 2012). One dataset was obtained from MADM forebrains in which all labeled cells are *wildtype* (*Emx:MADM:*+/+), while the two others were from MADM forebrains with heterozygous (*Emx:MADM:F*/+) or homozygous (*Emx:MADM:F/F*) deletions of *Egfr*. The *F*/+ and *F/F* MADM genotypes were phenotypically distinct with robust disruptions in production of glia, while largely preserving the neuronal production that developmentally occurs earlier than gliogenesis (Zhang et al., 2022).

In *Emx:MADM:F*/+ mice, red and green MADM cells correspond to *wildtype* and *Egfr-null* genotypes, respectively. While both red neurons and green neurons were present and relatively similar in density, a substantial increase in wild-type red MADM astrocytes and a visible absence of green *Egfr-null* MADM astrocytes was observable throughout the labeled regions of the forebrain (Figure 4A and 4B). In addition, no significant differences in total volume of the hemisphere, or regional volumes were observed, suggesting the MADM phenotypes were sparse enough to spare phenotypic changes in overall forebrain architecture. Figure 4C shows further region-wise comparison of red and green astrocyte densities between *Emx:MADM:*+/+ and *Emx:MADM:F*/+ brain hemispheres. Visualized in the left columns are percentage changes of regional cell densities while adjusted *p* values obtained from statistical tests are displayed on the right (see also Figure S2). Among 1326 ABA-annotated regions, 31 regions showed significant elevation of *wildtype* red astrocytes in the *Emx:MADM:F*/+ forebrain (Table S1).

**Figure 4.**
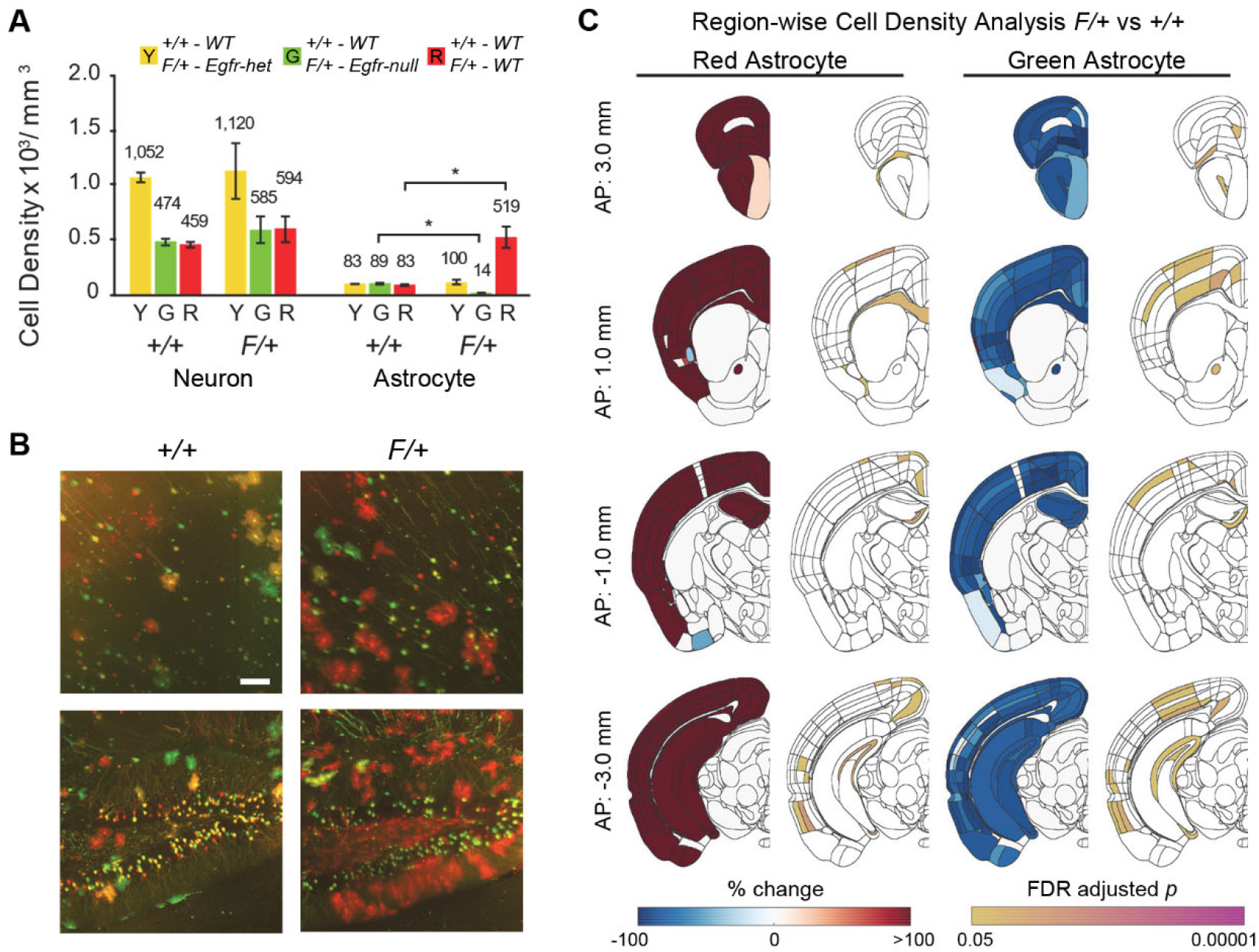
Elevated density of red MADM astrocytes and absence of green MADM astrocytes is faithfully captured by COMBINe in F/+ cortices. **(A)** Cell density comparison among between *Emx:MADM:*+/+ and *Emx:MADM:F*/+ mouse brain hemispheres (mean ± SD, n = 3; *, *p* < 0.05). In the *Emx:MADM:F*/+ forebrain, red and green cells correspond to *Egfr-null* and *wildtype* genotypes, respectively. While both red and green neurons were present, a substantial increase in wild type red astrocytes was revealed by COMBINe. **(B)** Representative images of the cortex and the hippocampus. Scale bar, 100 μm. **(C)** Region-wise analysis of average red and green astrocyte densities between *Emx:MADM:*+/+ and *Emx:MADM:F*/+ datasets. Left panels: percentage change in average cell densities of *Emx:MADM:F*/+ brain hemispheres compared to *Emx:MADM:*+/+ brain hemispheres. Right panels: adjusted *p*-values (n = 3). See also Figure S2 for analysis of other cell types. AP – anteroposterior distance taken from the bregma, FDR – false discovery rate.

### Using hierarchical analysis to discover major trends in regional densities of astrocytes

While statistical analysis of all ABA annotated forebrain regions (~1000) was useful, the average size of each brain region was relatively small. As such, the accuracy of registration to the ABA reference atlas is crucial to identify significantly altered densities of cell types, wherein small errors in the registration process can potentially mask significant results. Therefore, we sought to merge regions based on anatomical similarities. Toward this end we used the hierarchical structure of datasets in the ABA to merge regions and implement hierarchical analysis of region-wise statistics (Figure 5A, e.g., three-level hierarchical annotation of the somatomotor areas). Comparison of forebrain regions at multiple hierarchies revealed significant differences in red MADM astrocyte densities (Figure 5B; see also Table S2 for regions showing significant differences). Among the high-hierarchy regions, the corpus callosum showed significant increase of red astrocyte density in *Emx:MADM:F*/+ brain hemispheres compared to the control group (Figure 5C and 5D), which was consistent with manual observation of the acquired images.

**Figure 5.**
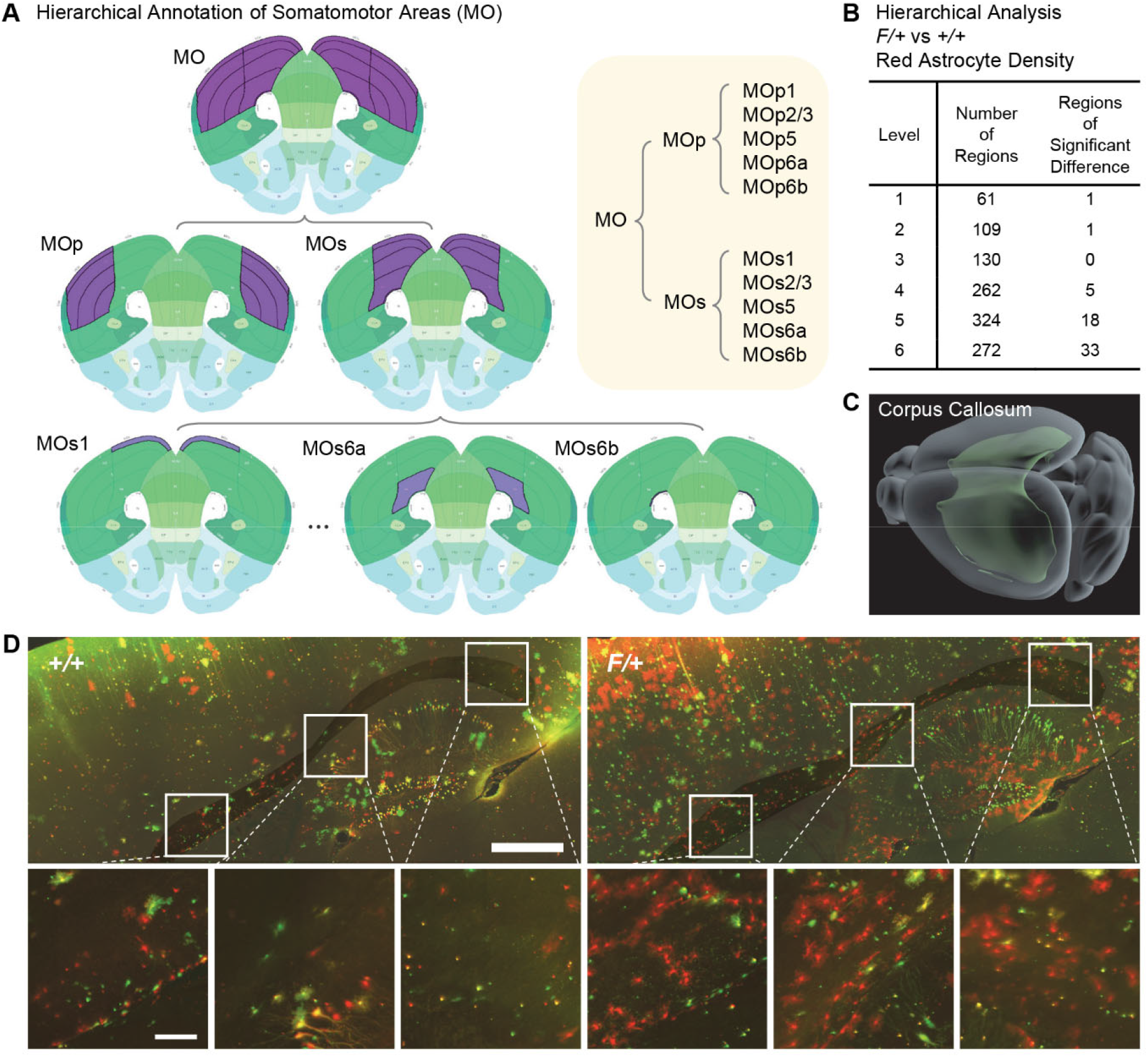
Hierarchical analysis reveals elevation in red MADM astrocytes in the *Emx:MADM:F*/+ corpus callosum. **(A)** Example of hierarchical annotations of somatomotor (MO) cortical areas according to the ABA. MOp – primary motor area. MOs – secondary motor area. **(B)** Results of hierarchical analysis revealed that the corpus callosum contains significantly elevated densities of red MADM astrocytes at hierarchy level 1. **(C)** 3D structure of the corpus callosum provided by ABA. **(D)** Representative images of the corpus callosum (scale bar, 500 μm) and zoomed in regions (scale bar, 100 μm).

### Variance in astrocyte distribution between rostral and caudal Emx:MADM:F/F cortices is quantitatively captured using COMBINe

Analysis of neurons and astrocytes in the *Emx:MADM:F/F* forebrain hemispheres using COMBINe confirmed a phenotype analyzed in our recent publication in rostral-caudal differences in EGFR-dependent and EGFR-independent astrocyte production (Zhang et al., 2022). It is important to note that all cells in the *Emx:MADM:F/F* forebrain are *Egfr-null* regardless of their MADM color. Figure 6A shows the average region-wise cell density maps onto the *Emx:MADM:F/F* isocortex as defined by the ABA. Hence, we compared the average astrocyte densities of selected regions and found that there was a significant difference in astrocyte distribution between rostral and caudal regions (Figure 6B). While astrocyte densities of control *Emx:MADM:*+/+ and *Emx:MADM:F/F* brain hemispheres appeared similar in caudal regions, the rostral regions of *Emx:MADM:F/F* cortices contained few, if any, astrocytes (Figure 6C). As expected, neurons that in general do not express EGFR showed similar distributions in the *Emx:MADM:F/F* and *Emx:MADM:*+/+ (Figure S3).

**Figure 6.**
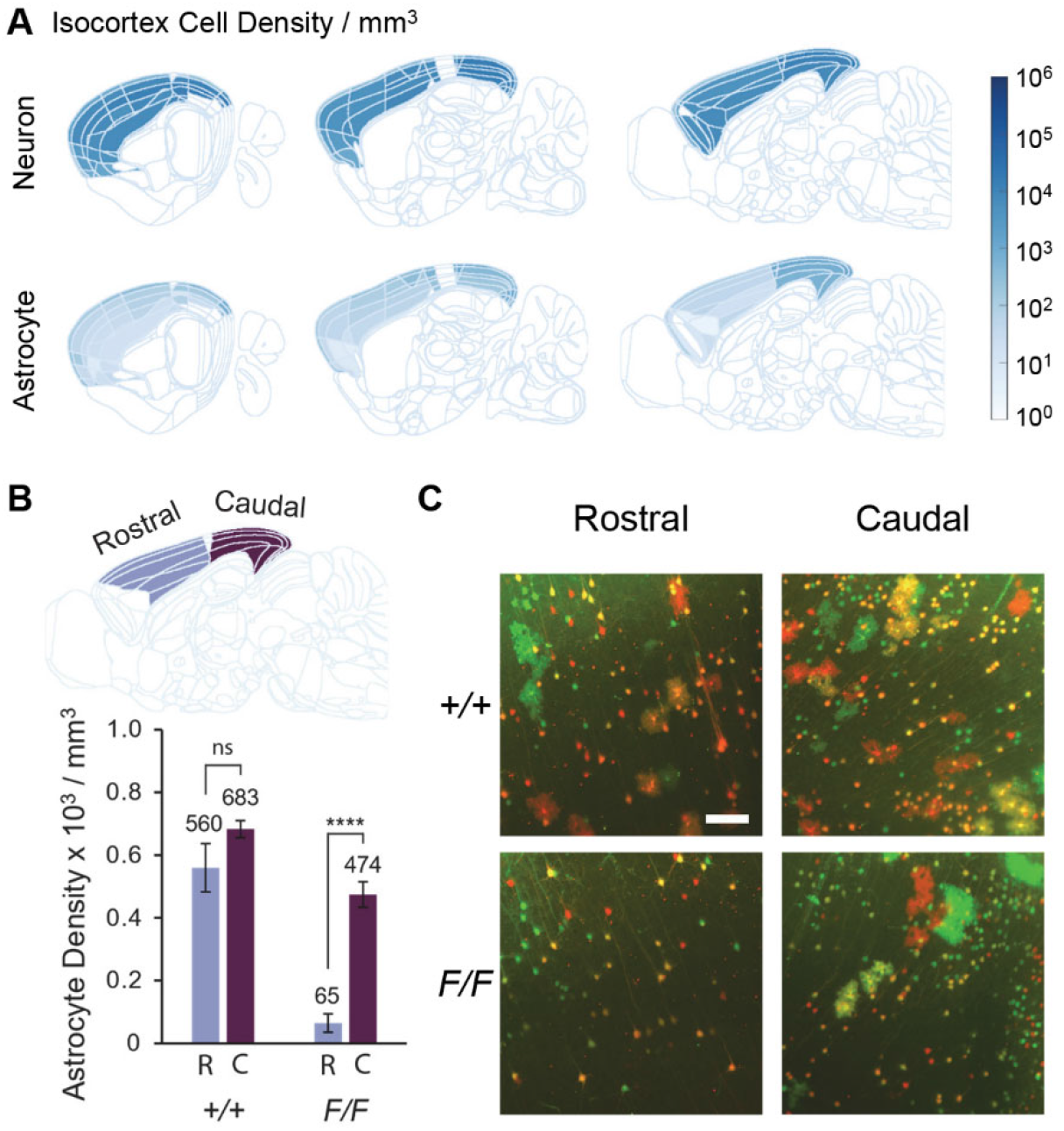
Variance in astrocyte distribution between rostral and caudal cortices is quantitatively captured using COMBINe. **(A)** Average regional cell density maps of *Emx:MADM:F/F* brain hemispheres (n = 3). **(B)** Significant difference in MADM-labeled astrocyte densities was observed between rostral and caudal regions in *Emx:MADM:F/F* brain hemispheres (bar chart: mean ± SD, n = 3; ****, *p* < 0.0001). See also Figure S3 for analysis of neuron densities. **(C)** Representative images of rostral and caudal areas (scale bars, 100 μm). Almost no MADM-labeled astrocytes are observable in rostral areas of *Emx:MADM:F/F* brain hemispheres.

To summarize, we compared the cellular populations’ densities among distinct *Emx1^cre^* induced MADM brain hemispheres with conditional deletion of *Egfr*. Region-wise analysis after registering the samples to ABA guided us to devise a hierarchical analysis method to characterize the variances observed in different cell populations among distinct genotypes. The revealed patterns matched laborious manual counting results which strongly validates our new COMBINe approach.

## Discussion

In this manuscript, we present COMBINe, an automated framework that can be applied to 3D imaging datasets that are captured with multiple color channels. To demonstrate the applicability of COMBINe, we image tissue cleared MADM-labeled mouse forebrains with cellular resolution. We have tested the influence of out-of-focus artifacts on the cell classification accuracy and show that if the illumination beams are parallel the RetinaNet performs adequately. COMBINe uses minimal storage space, as the datasets are not repeatedly saved per pipeline stage (e.g., after stitching and registration with the Allen Brain Atlas). We accomplish that by applying a bookkeeping strategy that uses the calculated transformation matrices in the stitching and registration parts to move the detected cells’ coordinates from one representation to the other. Consequently, only the raw tiles and a low-resolution stitched volume are required to complete the analysis. We also introduce a complementary method to perform the statistical analysis of our results, by comparing the cell densities on the hierarchical structure of the brain, provided by the ABA annotation. This method allows to average the cell densities across larger brain areas, that are morphologically and functionally connected, and to minimize the penalty for multiple comparisons. COMBINe can be adapted for other neuroscience studies that require quantitation of elements in the mouse brain using tissue clearing and immunofluorescence labeling. By enriching the framework components (e.g., adding features of segmentation and more) COMBINe can be applied to diverse neuroscience questions in need of analyzing 3D mouse brain datasets.

To test the RetinaNet model detection and classification success we have used the average precision measure, which is a widely used evaluation metric for object detection tasks (Everingham et al., 2015). The inference model (specific RetinaNet model) that is employed in the workflow achieved an average precision of 0.86 across six classes, showing its efficacy in cell detection and classification. However, when applied to large datasets, sample-wise quality control and common sense is necessary when utilizing learning-based methods. For example, in this study, the color ratio of neurons is referred to as an internal control to assess the cell detection process. Additionally, in the control brains, assessment that the color ratio is maintained across all tiles is used to assess outlier tiles.

Although we report an average precision of only 86%, such evaluation is performed based on human-annotated ground truth. Since there is a considerable variability even between human annotators, there are no perfect models for automated analysis. Hence, it is crucial to keep the analysis process consistent and eliminate potential biases.

Most of the data processing procedures in the current study were performed in 2D given the large size of the datasets and the differences in resolution along axes. In comparison with 3D processing, 2D processing has been intensely investigated and is applicable to diverse image data. Also, the labeling and manipulation of data in 2D is easier and more intuitive than in 3D. However, there are situations where 3D processing is favored. For example, when segmenting and tracing nerves, the structural information is mostly maintained in 3D. To summarize, processing in 2D or 3D should be carefully chosen given specific needs in analysis.

All in all, performance of COMBINe in analysis of *Emx1^cre^* MADM forebrains is in line with results obtained by manual counting in several ways; 1) restricted distribution of MADM cells, 2) quantitative differences between neurons and astrocytes, and 3) the distribution of different cell colors across the brain. We did not observe volumetric changes when the experimental groups were compared with the control, confirming that sparse alterations using MADM which is estimated to occur in 1:1000 cells in the brain (Hippenmeyer et al., 2010; Zong et al., 2005), has undetectable effects on overall tissue development and homeostasis.

From the biological perspective, COMBINe quantitatively and unbiasedly confirms two important phenotypes in the three *Egfr* genetic backgrounds described here. The findings from *Emx:MADM:F*/+ forebrains suggest that when faced with a largely *Egfr-heterozygous* forebrain background, sparse populations of red *wildtype* MADM cells may overexpand to generate a larger number of astrocytes than their normal capacity during perinatal development when gliogenesis occurs. In contrast, *Egfr-null* MADM cells fail to generate astrocytes in the rostral part of the isocortex. We found similar results in our recent clonal study using tamoxifen-inducible *Nes-creER:MADM:F*/+ alleles to label gliogenic progeny at clonal densities, and found a robust dosage response to EGFR expression in the generation of both astrocytes and oligodendrocytes (Zhang et al., 2020). Since the *wildtype* glial progenitors are essentially overexpressing EGFR relative to their surrounding cells in the *Emx:MADM:F*/+ background, these results constitute a gain-of-function scenario that explains the strong effect on astrocyte production. An unexpected result in our current study was the region-restricted presence of glia in *Emx:MADM:F/F* forebrains, since the F/+ findings suggested that no astrocytes should be produced in the *homozygous Egfr-null* background. However, the astrocytes in this case were largely in the caudal, but not rostral forebrain regions. This suggests the possibility of an EGFR-independent gliogenic progenitor pool in the caudal aspect of the forebrain, appears capable of compensating for the loss of EGFR-dependent glia which are critical for astrocyte production in rostral forebrain regions. Interestingly, the EGFR-dependent astrocytes appear to occupy neocortical and possibly other neo-forebrain areas, whereas the EGFR-independent populations which are present in the *Emx:MADM:F/F* appear to seed evolutionary older paleocortical and hippocampal areas. The precise identity and anatomical domain of the EGFR-independent glia producing progenitors remains to be determined.

## Limitation of Study

Currently, deep learning is still considered a black box i.e., not explainable, which makes it difficult to evaluate its failures and improve the model performance (Belle & Papantonis, 2021). Moreover, as deep learning is a data-driven method, additional training is required before applying it to a different sample type. Furthermore, the packages utilized/modified in this study were built on different operating systems (e.g., Linux and Windows) and modifying them to run in a different operating system is challenging and inconvenient to use. In addition, performance of the brain registration approach adopted in this pipeline will drop if the sample is incomplete. Last, some of the steps still require manual check-ins to ensure proper functioning of COMBINe. For example, it is highly recommended to manually inspect a few images in each tile after channel alignment to make sure the alignment is accurate, and the cell detection model needs to be validated every time when analyzing a new batch of datasets.

## Acknowledgements

The authors would like to thank the Greenbaum lab members for discussions and assistance in revising the manuscript. Grants to HTG and AG (R21NS129093 and R56NS117019) supported this work.

## Author Contributions

Conceptualization: YC, AG, HTG; Methodology: YC, CL; Investigation: YC; Software: YC; Visualization: YC; Resources: XZ, HTG; Supervision: AG, HTG; Writing – original draft: YC, XZ; Writing – review & editing: YC, AG, HTG

## Declaration of Interests

The authors declare no competing interests.

## Materials and methods

### Animals

Mice were housed in a 12-hour light:dark cycle with ad libitum access to food and water. All procedures were performed under the regulations and approval from Institutional Animal Care and Use Committee and at North Carolina State University. *Emx1^cre^:MADM11^TG/GT^:Egfr*^+/+^, *Egfr*^*F*/+^ and *Egfr^F/F^* mice were generated using breeding schemes previously described (Zhang et al., 2020). In this study, three brain hemispheres collected from three mice of each genotype were tissue cleared and analyzed.

### Tissue clearing

The iDISCO+ protocol (Renier et al., 2016) was used to stain and clear the mouse brain hemispheres. Brain hemispheres were dehydrated at room temperature (RT) using methanol gradients diluted in ddH_2_O: 20%, 40%, 60%, 80%, 100%, 100%; 1 h each. Brain hemispheres were then chilled at 4 °C for 2 h and submerged in the mixture of dichloromethane and methanol (v:v=2:1) overnight at RT. After washing twice in 100% methanol at RT, brain hemispheres were chilled at 4 °C for 2 h and decolorized in chilled fresh 5% H_2_O_2_ in methanol overnight at 4 °C. After bleaching, brain hemispheres were rehydrated with methanol series diluted in ddH_2_O (100%, 80%, 60%, 40%, 20%), followed by one wash with PBS and two washes with PBS with 0.02% Triton X-100, 1 h each at RT. Next, brain hemispheres were permeabilized with permeabilization solution (0.2% Triton X-100, 0.3M Glycine, 20% DMSO in PBS) for 2 d at 37 °C and blocked with blocking solution (0.2% Triton X-100, 6% donkey serum, 10% DMSO in PBS) for 2 d at 37 °C. Brain hemispheres were then incubated with chicken anti-GFP (1:200, Aves Labs) and rabbit anti-RFP (1:200, Rockland) in PTwH (0.2% Tween-20, 10 mg/L heparin in PBS) with 5% DMSO (v/v) and 3% donkey serum for 7 d at 37 °C. After primary staining, brain hemispheres were washed 5 times with PTwH, 2 h each until the next day. Next, brain hemispheres were incubated with donkey anti-chicken AlexaFluor 647 (1:200, Jackson ImmunoResearch) and donkey anti-rabbit Cy3 (1:200, Jackson ImmunoResearch) in PTwH with 3% donkey serum for 5 d at 37 °C, followed by 5 washes with PTwH. For optical clearing, brain hemispheres were dehydrated with methanol series diluted in ddH2O (20%, 40%, 60%, 80%, 100%), 1 h each at RT, and incubated in the mixture of dichloromethane and methanol (v:v=2:1) for 3 h at RT. Brain hemispheres were then submerged in dichloromethane 15 minutes twice at RT. All the incubation steps above were performed with gentle shaking. Brain hemispheres were cleared with dibenzyl ether at RT.

### Light-sheet imaging

Cleared mouse brain hemispheres were imaged using a custom-built light sheet fluorescence microscope, of which the setup was outlined in (Li et al., 2021, 2022; Moatti et al., 2020). Each brain hemisphere was imaged by illumination of 561 nm and 640 nm respectively, with a voxel size of 0.65 × 0.65 × 10 μm^3^. During image acquisition, every 1 mm in z direction, autofocus method would be applied to correct for the drift between the light-sheet plane and the detection focal plane.

### Channel alignment

When the illumination beams of different wavelengths are parallel, images of different color channels can be matched through translation in x, y, and z-axes. NuMorph (Krupa et al., 2021) was employed for local channel alignment on each imaged tile (MATLAB). First, displacement in z-planes was determined by maximizing phase correlation between two channels. After adjustment in z, color registration was performed in x and y-axes.

### Cell detection

Both the training data and the validation data contained fluorescence images from different imaging modalities, e.g., slide scanner, confocal fluorescence microscopy, and LSFM (Y. Cai et al., 2021). 3361 individual cells (2497 neurons and 864 astrocytes) were labeled in the training data. 1164 individual cells (896 neurons and 268 astrocytes) were labeled in the validation data. Test data was generated from light sheet datasets that were analyzed in this paper. 1664 individual cells (1462 neurons and 202 astrocytes) were labeled in the test data.

A RetinaNet (Lin et al., 2018) model was trained to detect cells of six classes: yellow, red, green neurons and astrocytes. The repository cloned from the source (https://github.com/fizyr/keras-retinanet) was adapted to this work (https://github.com/yccc12/keras-retinanet). Transfer learning was adopted with a pre-trained ResNet50 being used as the backbone. Data augmentation strategies such as geometrical transforms, color swap and saturation simulation were applied (Y. Cai et al., 2021). The initial learning rate was 0.0001 and the batch size was four. Using an Adam optimizer, the model was trained for 50 epochs. The model with the best performance on validation data was selected as the inference model.

Because stitching was likely to generate artifacts, cell detection was performed per image tile, during which images were cropped into 512 × 512 image patches with an overlap fraction of 0.125 for inference. Detections in overlapped areas were merged based on intersection over union and confidence score.

### Stitching

TeraStitcher (Bria & Iannello, 2012) advanced mode was used to stitch the acquired datasets, with misalignment across tiles being corrected. Parameters output from TeraStitcher were adopted to stitch detected cells of each tile. Duplicate detections in overlapping regions between adjacent tiles were rejected. Redundant detections across different z-planes were merged based on signal intensities when the output bounding boxes between adjacent z-planes were found overlapped, which was determined by an intersection over union above 0.5 between two bounding boxes. Stitched datasets and detected cells were visualized in Imaris 9.5 (Oxford Instruments).

### Registration

ClearMap (Renier et al., 2016) in Linux was utilized to register stitched datasets to Allen Brain Atlas (Wang et al., 2020) and to map detected cells to annotated brain regions

### Statistical analysis

Regional cell densities between different groups were compared in MATLAB using unpaired t-tests assuming unequal variances. *p*-values were adjusted by the Benjamini–Hochberg procedure to control the false discovery rate (< 0.05) for simultaneous multiple testing.

### Visualization

Cell densities and statistics were visualized in MATLAB using a script modified from NuMorph.

## Notes

### Competing Interest Statement

The authors have declared no competing interest.

