## Supplemental Figures and Tables for "COMBINe: Automated Detection and Classification of Neurons and Astrocytes in Tissue Cleared Mouse Brains"

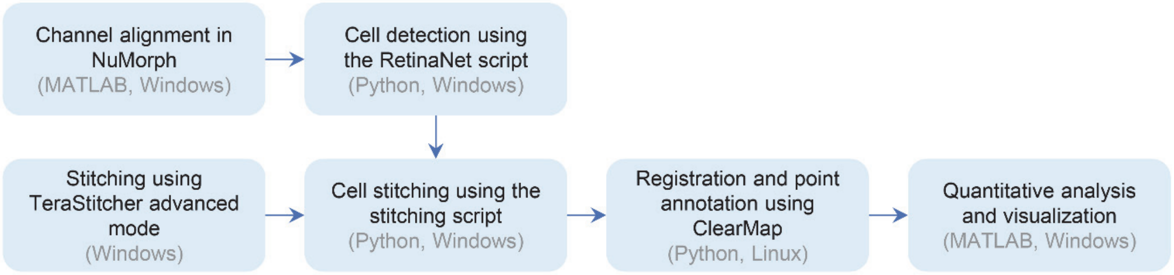


**Figure S1. Running the pipeline for MADM cell mapping.**

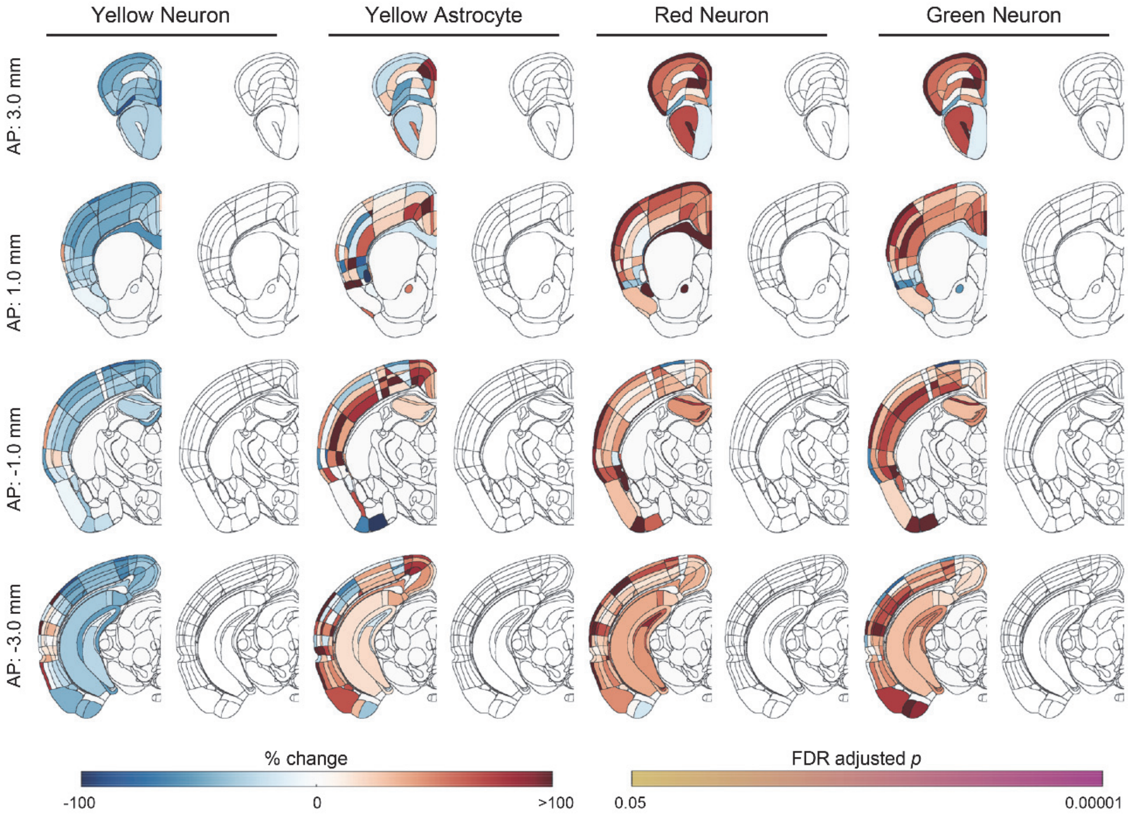


**Figure S2. Region-wise analysis of average cell densities between *Emx:MADM:+/+* and *Emx:MADM:F/+* datasets.** Left panels: percentage change in average cell densities of *Emx:MADM:F/+* brain hemispheres compared to *Emx:MADM:+/+* brain hemispheres. Right panels: adjusted *p*-values (n = 3). AP – anteroposterior distance taken from the bregma, FDR – false discovery rate.

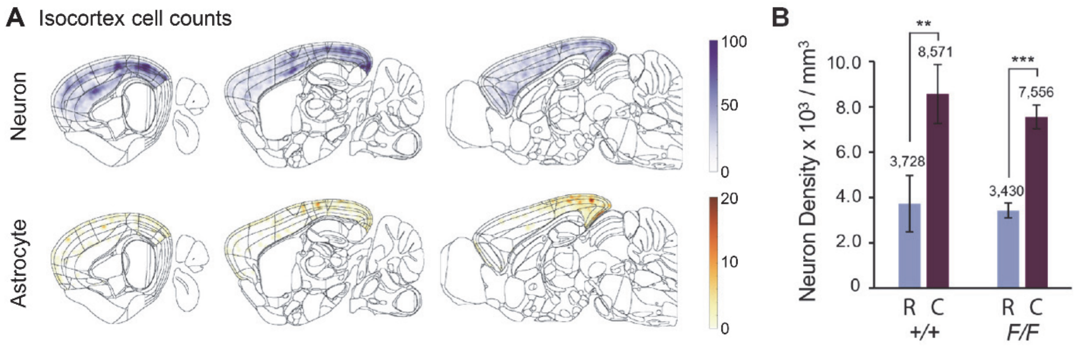


**Figure S3. Variance in neuron distribution between rostral and caudal cortices. (A)** Average voxel-wise cell density maps of *Emx:MADM:F/F* brain hemispheres (n = 3). **(B)** Significant difference in MADM-labeled neuron densities was observed between rostral and caudal regions in both *Emx:MADM:+/+* and *Emx:MADM:F/F* brain hemispheres (bar chart: mean ± SD, n = 3; **, $p < 0.01$; ***, $p < 0.001$).

Table S1. 31 regions show significant increase of red MADM astrocyte.

| Name | % Change | Adjusted *p*-value |
|---|---|---|
| Cortical amygdalar area | 2379.618 | 0.0027 |
| Cortical amygdalar area posterior part | 2530.341 | 0.0027 |
| Primary motor area Layer 1 | 659.7982 | 0.0038 |
| Entorhinal area lateral part layer 3 | 418.6971 | 0.0038 |
| Corpus callosum | 469.5149 | 0.0074 |
| Dentate gyrus granule cell layer | 1265.942 | 0.0107 |
| Orbital area | 328.3208 | 0.0121 |
| Retrosplenial area lateral agranular part layer 2/3 | 247.3442 | 0.0182 |
| Genu of corpus callosum | 438.9664 | 0.0221 |
| Entorhinal area medial part dorsal zone layer 5 | 445.1044 | 0.024 |
| Retrosplenial area dorsal part | 515.8460 | 0.0261 |
| Anteromedial visual area layer 5 | 627.4307 | 0.0265 |
| Agranular insular area | 501.1534 | 0.0274 |
| Supplemental somatosensory area layer 6b | 306.5585 | 0.0309 |
| Ventral auditory area layer 5 | 1369.305 | 0.0419 |
| Anteromedial visual area | 338.6888 | 0.0419 |
| Posteromedial visual area layer 1 | 937.5416 | 0.0419 |
| Retrosplenial area | 425.3429 | 0.0419 |
| Retrosplenial area dorsal part layer 1 | 372.9179 | 0.0419 |
| Retrosplenial area ventral part | 395.9375 | 0.0419 |
| Entorhinal area lateral part | 442.4262 | 0.0419 |
| Entorhinal area lateral part layer 2 | 383.3483 | 0.0419 |
| Lateral forebrain bundle system | 435.2953 | 0.0419 |
| Retrosplenial area ventral part layer 5 | 504.5898 | 0.0438 |
| Temporal association areas layer 6a | 825.1289 | 0.0450 |
| Entorhinal area medial part dorsal zone layer 3 | 308.5111 | 0.0450 |
| Primary somatosensory area upper limb layer 1 | 328.5355 | 0.0454 |
| Triangular nucleus of septum | 538.6442 | 0.0454 |
| Olfactory areas | 250.8286 | 0.0475 |
| Anterior cingulate area dorsal part | 599.8512 | 0.0499 |
| Infralimbic area layer 6a | 609.0158 | 0.0499 |

Table S2. Regions showing significant differences in hierarchical analysis.

| Level | Name |
|---|---|
| 1 | Corpus callosum |
| 2 | Genu of corpus callosum |
| 3 | - |
| 4 | Cortical amygdalar area<br>Orbital area<br>Agranular insular area<br>Retrosplenial area<br>Triangular nucleus of septum |
| 5 | Cortical amygdalar area posterior part<br>Secondary motor area<br>Gustatory areas layer 5<br>Visceral area layer 4<br>Anteromedial visual area<br>Posterolateral visual area<br>Anterior cingulate area dorsal part<br>Infralimbic area layer 5<br>Infralimbic area layer 6a<br>Infralimbic area layer 6b<br>Retrosplenial area lateral agranular part<br>Retrosplenial area dorsal part<br>Retrosplenial area ventral part<br>Temporal association areas layer 5<br>Temporal association areas layer 6a<br>Dentate gyrus<br>Entorhinal area<br>Prosubiculum |
| 6 | Primary motor area Layer 5<br>Secondary motor area layer 1<br>Secondary motor area layer 2/3<br>Secondary motor area layer 5<br>Supplemental somatosensory area layer 6b<br>Dorsal auditory area layer 4<br>Ventral auditory area layer 5<br>Ventral auditory area layer 6b<br>Anteromedial visual area layer 1<br>Anteromedial visual area layer 5<br>Anteromedial visual area layer 6a<br>Anteromedial visual area layer 6b<br>Lateral visual area layer 4<br>Primary visual area layer 1<br>Posterolateral visual area layer 2/3<br>Posterolateral visual area layer 4<br>Posteromedial visual area layer 1<br>Posteromedial visual area layer 2/3<br>Anterior cingulate area dorsal part layer 5<br>Anterior cingulate area dorsal part layer 6a<br>Agranular insular area dorsal part layer 2/3 |

| | |
|---|---|
| | Retrosplenial area lateral agranular part layer 1<br>Retrosplenial area lateral agranular part layer 2/3<br>Retrosplenial area lateral agranular part layer 6b<br>Retrosplenial area dorsal part layer 1<br>Retrosplenial area dorsal part layer 2/3<br>Retrosplenial area dorsal part layer 5<br>Retrosplenial area ventral part layer 5<br>Cortical amygdalar area posterior part medial zone<br>Dentate gyrus molecular layer<br>Dentate gyrus polymorph layer<br>Entorhinal area lateral part<br>Entorhinal area medial part dorsal zone |
